# The last common ancestor of most bilaterian animals possessed at least 9 opsins

**DOI:** 10.1101/052902

**Authors:** MD Ramirez, AN Pairett, MS Pankey, JM Serb, DI Speiser, AJ Swafford, TH Oakley

**Affiliations:** Department of Ecology, Evolution and Marine Biology, University of California, Santa Barbara; Department of Ecology, Evolution and Organismal Biology, Iowa State University; Department of Molecular, Cellular and Biomedical Sciences, University of New Hampshire; Department of Biological Sciences, University of South Carolina, Columbia, SC

**Keywords:** reconciled tree, eye evolution, extraocular photoreceptors, phototransduction, vision

## Abstract

The opsin gene family encodes key proteins animals use to sense light and has expanded dramatically since it originated early in animal evolution. Understanding the origins of opsin diversity can offer clues to how separate lineages of animals have repurposed different opsin paralogs for different light-detecting functions. However, the more we look for opsins outside of eyes and from additional animal phyla, the more opsins we uncover, suggesting we still do not know the true extent of opsin diversity, nor the ancestry of opsin diversity in animals. To estimate the number of opsin paralogs present in both the last common ancestor of the Nephrozoa (bilaterians excluding Xenoacoelomorpha), and the ancestor of Cnidaria + Bilateria, we reconstructed a reconciled opsin phylogeny using sequences from 14 animal phyla, especially the traditionally poorly-sampled echinoderms and molluscs. Our analysis strongly supports a repertoire of at least nine opsin paralogs in the bilaterian ancestor and at least four opsin paralogs in the last common ancestor of Cnidaria + Bilateria. Thus, the kernels of extant opsin diversity arose much earlier in animal history than previously known. Further, opsins likely duplicated and were lost many times, with different lineages of animals maintaining different repertoires of opsin paralogs. This phylogenetic information can inform hypotheses about the functions of different opsin paralogs and be used to understand how and when opsins were incorporated into complex traits like eyes and extraocular sensors.

## Introduction

As the protein component of visual pigments, opsins are used in the majority of light-detecting cells found in animals (Nilsson 2013; Oakley & Speiser 2015). Opsins are G-protein coupled receptors which bind a light-sensitive chromophore via a Schiff base linkage at a conserved lysine residue (Terakita 2005). When the chromophore absorbs a photon, conformational changes in the chromophore and opsin protein activate G-protein signal transduction (Terakita 2005). Despite their widespread importance in animal photosensitivity, most work on the function and evolution of opsins focused initially on those expressed in the eyes of vertebrates and arthropods (Nathans & Hogness 1983; O’Tousa et al. 1985). Only more recently has work on opsins included those expressed outside eyes or from other animal phyla (Velarde et al. 2005; Radu et al. 2008; Hering et al. 2012; D’Aniello et al. 2015; Hering & Mayer 2014). We now know the evolutionary history of opsins is one of many gains and losses of genes across time and among species (Colbourne et al. 2011; Henze & Oakley 2015; Davies et al. 2015; Liegertová et al. 2015; Feuda et al. 2016). Understanding this kind of high gene turnover requires broad taxonomic sampling of opsins to fully reconstruct their evolutionary origins, especially because we know that ancient losses may result in the complete absence of some opsin paralogs, even in major groups of animals. Previous large-scale opsin phylogenies have also found many sequences that fall outside of the well-known opsin groups, typically identified in taxa for which we have sparse data, e.g. arthropsins in *Daphnia* or Echinopsins B in echinoderms (e.g. Colbourne et al. 2011; D’Aniello et al. 2015). Most analyses do not address the nature of these orphaned sequences. While they may be recently-diverged, lineage-specific duplications, another possibility is that they represent entire opsin paralogs that are absent from the phyla that have been most heavily sampled. Without an accurate picture of how opsin paralogs are distributed among animals, it is challenging to address how diverse opsins really are, when that diversity arose, and when and how different opsins were integrated into different kinds of light-detecting structures during evolution.

Opsins evolved very early in animals (Plachetzki et al. 2007; Feuda et al. 2012; Oakley & Speiser 2015), likely first expressed in light-sensitive cells and later in more complex structures like eyes (Arendt & Wittbrodt 2001; Nilsson 2013). Historically, opsin diversity has been partitioned among three clades: the ‘ciliary’ or c-opsins, the ‘rhabdomeric’ or r-opsins, and ‘Group 4 opsins’ *sensu* (Porter et al. 2012; Liegertová et al. 2015). We propose renaming the c-and r-opsin paralogs ‘canonical c-opsin’ and ‘canonical r-opsin’ to denote the originally described visual opsins of vertebrates and invertebrates (Terakita 2005). We also propose renaming the ‘Group 4 opsins’ coined by Porter et al. (2012) as tetraopsins. Hereafter, we refer to these opsins by these new names. A possible fourth clade of opsins, cnidops, are currently known only from cnidarians and include sequences from all major cnidarian classes (Plachetzki et al. 2007; Feuda et al. 2012). To understand how many opsin paralogs were present in the last common eumetazoan and bilaterian ancestors, we need to understand when these major opsin clades arose and how they are related to each other.

Because cnidarians represent a rare non-bilaterian animal lineage with opsins, their opsin repertoire provides key insights into the opsin paralogs present in the last common ancestor of eumetazoans. However, relating cnidarian opsins to the major animal opsin paralogs has proved difficult, and hypotheses on how cnidarian and bilaterian opsins relate vary widely between analyses. For example, some analyses indicate cnidarian genomes encode either cnidops alone (Porter et al. 2012; Liegertová et al. 2015), or cnidops plus c-opsins (Plachetzki et al. 2007; Vopalensky & Kozmik 2009). However, others suggest the most recent ancestor of eumetazoans had three opsin paralogs: c-opsins, r-opsins and tetraopsins (Suga et al. 2008; Feuda et al. 2012; Bielecki et al. 2014; Feuda et al. 2014). Based on *in-vitro* assays, an opsin from the coral *Acropora palmata* (Acropsin3) interacts with the same G-protein q alpha subunit used by r-opsins (Lee et al. 1994; Mason et al. 2012). Together with the hypothesized phylogenetic position of this opsin, the functional test suggests that some cnidarians may possess canonical r-opsins (Mason et al. 2012). Still, the exact placement of this and other cnidarian opsins is highly sensitive to the specific substitution models and gene sampling regime used in each analysis.

Reconstructing opsin evolution in bilaterians poses yet more challenges. Early estimates of opsin diversity in the last common bilaterian ancestor identified two (Nilsson 2005) or three (Plachetzki et al. 2007; Porter et al. 2012; Feuda et al. 2012, 2014) paralogs, corresponding to the canonical c-opsins and canonical r-opsins, or canonical c, canonical r, and tetraopsins, respectively. Recent sampling efforts to survey new taxa and extraocular tissues have expanded our current view of opsin diversity, and we now recognize that multiple clades of opsins found in extant animals were present in the last common ancestor of bilaterians, based on their presence in both deuterostome (e.g. vertebrates and echinoderms) and protostome (e.g. arthropods and molluscs) genomes. These considerations indicate at least five opsin paralogs in the last common ancestor of bilaterians (Terakita 2005; Vopalensky & Kozmik 2009; Suga et al. 2008) or six (Hering & Mayer 2014; Liegertová et al. 2015; Feuda et al. 2014), distributed between the bilaterian c-, r-and tetraopsins. With these additions, a pattern emerges: as we catalog opsins in diverse phyla and from different types of light receptors, we uncover a greater diversity of opsin paralogs. A further wrinkle is recent strong support for the hypothesis that Acoelomorpha and Xenoturbella together are sister (as Xenacoelomorpha) to other bilaterians (Cannon et al. 2016), rather than nested within the deuterostomes. Thus all estimates of the number of opsin paralogs in the last common bilaterian ancestor need to include Xenacoelomorpha opsins, which are missing from all opsin phylogenies to date (including our analysis here). While estimates may hold once Acoelomorpha are considered, to be conservative, at present we can only truly infer the opsin repertoire for Nephrozoa, and the last common ancestor of protostomes (excluding Xenacoelomorpha) and deuterostomes.

A primary goal of our analysis is to reconstruct a more taxonomically comprehensive evolutionary history of animal opsins to understand the origins of bilaterian opsin diversity. We achieve this in two ways. First, we include newly published opsin sequences from multiple studies that have yet to be synthesized in a large scale phylogenetic analysis. Second, we identify additional new opsins from both publicly available transcriptomes and nine unpublished mollusc transcriptomes, as molluscs are the second most speciose phylum but lag far behind other large taxa in terms of representation in opsin phylogenies to date. With this more comprehensive data set, we produced the first large-scale formally reconciled opsin phylogeny and we use it to more explicitly estimate the number of opsins present in the last common ancestor of Protostomia + Deuterostomia. This approach allows us to infer at least nine opsin paralogs were likely present in early bilaterians. Further, from the distribution of cnidarian opsins, we infer that the last common ancestor of eumetazoans had at least four opsin paralogs. These results suggest a radiation in opsin diversity prior to the origin of bilaterians, followed by unique patterns of duplications and losses specific to different animal lineages. Finally, these results urge a renewed focus on surveying opsins in understudied phyla (prime candidates include Annelida and Xenacoelomorpha, and non-bilaterians like Cnidaria and Ctenophora), and learning the functions of those opsins.

## Methods

### Data collection

We searched both NCBI and UniProt using BLAST (Gish & States 1993) with a bait set of 5 opsin sequences (accession numbers: BAG80696.1; NP_001014890.1; CAA49906.1; O15974.1; P23820.1) and an e-value cutoff of 1e-5. Our goal was at first to maximize the identification of potential opsins from understudied taxa, so we excluded vertebrates and arthropods from our BLAST search on NCBI and downloaded the top 250 hits per opsin bait. We then searched Uniref90 with the same bait sequences and cutoff value, then downloaded only lophotrochozoan (NCBI taxonomic ID: 1206795) sequences/clusters. We combined all the sequences we recovered from NCBI and Uniref90 with sequences from other publications, which include tardigrades, arthropods, ambulacraria, cubozoan cnidarians and vertebrates (Hering & Mayer 2014; Henze & Oakley 2015; D’Aniello et al. 2015; Liegertová et al. 2015; Davies et al. 2015). To this initial database of published sequences, we added mollusc opsins that we gathered by running Phylogenetically Informed Annotation, PIA, (Speiser et al. 2014) on transcriptomes and NCBI TSAs from 2 cephalopods, 3 chitons, 5 gastropods, and 3 bivalves.

### Data grooming

Because our initial data collection was permissive, our raw dataset (over 1,600 sequences) contained duplicate sequences as well as a number of non-opsin GPCRs. We used CD-HIT (Li & Godzik 2006; Fu et al. 2012) to cluster together sequences that were more than 90% similar, allowing us to remove duplicates and highly similar sequences. To remove non-opsin GPCRs, we first ran the dataset through SATé-II (Liu et al. 2012). SATé-II employs FastTree 2 (Price et al. 2010) on an initial MAFFT (Katoh & Standley 2013) alignment before subdividing the alignment into 200 subproblems and realigning with MAFFT. The realigned subproblems are then merged using MUSCLE (Edgar 2004), and a new tree produced by FastTree, and the maximum likelihood (ML) score is calculated. SATé-II iterates until a pre-defined stopping point or likelihood improvement. Next, we used FigTree (Rambaut 2007) to visualize trees, rooted with melatonin receptors (Feuda et al. 2014; Plachetzki et al. 2010). We then trimmed this tree to exclude non-opsins using a custom python script called Supercuts (Swafford 2016) and retained the ingroup clade for subsequent analyses. We removed any sequences from the alignment that lacked the conserved lysine residue homologous to K296 of bovine rhodopsin. We also manually trimmed the beginning and end of the alignment to the first and last aligned blocks using Aliview (Larsson 2014). Finally, although they lack the conserved lysine, we added the *Trichoplax adherens* placopsins back to our dataset as a close outgroup to root our tree, as Feuda et al. (2012) showed that placopsins are sister to all other animal opsins. In total, our groomed dataset had 768 opsins with the conserved K296 (plus three placozoan opsins without the lysine) from 248 species across 14 phyla.

### Tree estimation and support values

To create the final alignment for our dataset, we ran SATé on our dataset using the following configuration: a subproblem fraction of 0.025, stopping iterations after 5 unimproved ML scores and FastTree under the GTR gamma model. We next used the MPI version of IQ-TREE 1.4.0 (Nguyen et al. 2014), to select a substitution model based on our SATé alignment, infer a maximum likelihood tree, and compute support values. IQ-TREE incorporates an approach for calculating ultrafast bootstraps (UFBoot), which may have fewer biases compared to other bootstrapping methods (Minh et al. 2013). We were also able to perform the SH-like approximate likelihood ratio test (SH-aLRT) and the approximate Bayes test as implemented in IQ-TREE to assess support for single branches to complement our UFBoot analysis (Guindon et al. 2010; Anisimova et al. 2011). SH-aLRT branch supports are often more consistent and conservative than bootstrapping methods (Simmons & Randle 2014; Simmons & Norton 2014). The IQ-TREE substitution model test selected the LG+F+R8 model for our alignment based on BIC. Because we had a large number of relatively short sequences, we performed 50 ML tree searches varying the perturbation value (0.1-0.5). IQ-TREE keeps the best ML tree while it searches the tree parameter space, and stops searching after going through a user-defined number of tree. We extended this number to 500 trees to better explore tree parameter space. Two trees had virtually identical high log-likelihood scores, and so we ran IQ-TREE again, setting each tree as the starting tree, to break the tie and to get UFBoot, SH-aLRT and aBayes values for the final, highest log-likelihood tree. The code used for this analysis, our dataset and the resultant tree are available on BitBucket (UCSB Phylogenetics).

### Tree reconciliation and rearrangement

We used NOTUNG 2.8 (Chen et al. 2000) to reconcile the gene tree with a metazoan species tree based on NCBI Taxonomy. This animal phylogeny places sponges as sister to all other animals, and posits unresolved relationships between ctenophores, cnidarians and bilaterians. While the order of branching in early metazoans is contentious, we do not expect it to affect our estimate of the number of paralogs present in the last common ancestor of Cnidaria + Bilateria. Because we were unable to include Xenoacoelomorpha opsins in our phylogeny, our estimate of bilaterian opsin paralogs applies only to the last common ancestor of protostomes and deuterostomes. To perform both a reconciliation and rearrangement of weakly supported branches, NOTUNG requires a fully resolved species tree. We used the ape package (Paradis et al. 2004) in R (R Core Team 2016) to randomly resolve polytomies present in the species tree. Because our analysis focuses on major splits in the animal phylogeny that are well supported, e.g. protostomes vs deuterostomes, the random resolution of more shallow nodes did not impact our results. We set the penalty for duplications to 1.5, losses to 1.0 and the edge weight threshold for rearrangement to 95.0 (i.e. nodes with >95% UFBoot support are rearranged to minimize duplication and loss costs).

### Tree visualization

We used FigTree 1.4.2 (Rambaut 2007) and TreeGraph2 (Stöver & Müller 2010) to collapse opsin clades by hand according to major taxonomic group (chordates, echinoderms, lophotrochozoans or ecdysozoans), and Evolview (Zhang et al. 2012) to format the tree color, branch length, etc. for Figure 1. We used iTOL (Letunic & Bork 2011) to combine the tallies of opsins per phylum or molluscan class with animal and mollusc phylogenies (Figures 4 and 5). For Supplemental Figures S1 and S2 we used ETE3 to build and annotate the trees (Huerta-Cepas et al. 2016). Data and code for these trees are also available at the UCSB Phylogenetics BitBucket. We made final adjustments to the outputs of these programs using OmniGraffle Pro (v. 6.5, Omni Group, Seattle, WA).

**Figure 1.**
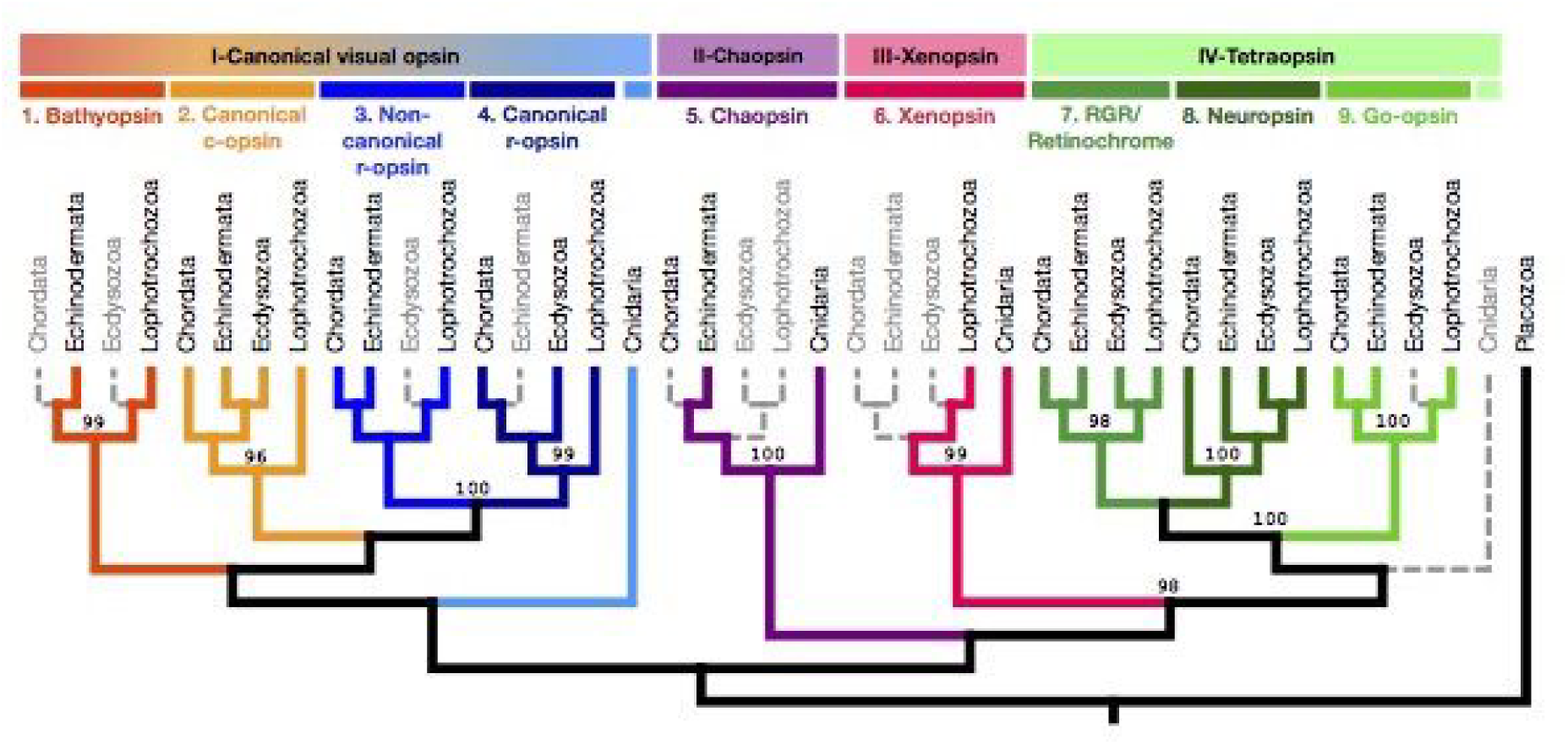
There are nine bilaterian opsin paralogs spread among four major eumetazoan opsin paralogs. The four major eumetazoan opsin paralogs are indicated at the top with roman numerals. The nine bilaterian opsin paralogs are indicated with arabic numerals and are color coded to match the corresponding branches. Each opsin clade has been reconciled and collapsed into four major taxonomic groups: chordates, echinoderms, ecdysozoans and lophotrochozoans. Colored branches indicate the presence of an opsin in at least one species within the major taxonomic group. Light gray dashed branches indicate absence of an opsin paralog from the taxa indicated at the tips. We infer these absences represent true losses of that opsin paralog. Ultrafast bootstrap (UFBoot) supports from IQ-TREE are given at the nodes they support. All unlabeled nodes had UFBoot supports below 95% and were rearranged during tree reconciliation. See Suppl. Figure S2 for the uncollapsed reconciled tree.

## Results

From our reconciled tree containing 768 unique sequences, our analysis strongly supports at least nine bilaterian opsin paralogs spread across four eumetazoan paralogs (Figures 1 & 2, complete gene and reconciled gene trees in Suppl. Figures S1 & S2). We recover the six bilaterian opsin paralog groups described in previous publications: canonical c-opsin, canonical r-opsin, Go-opsin, RGR/retinochrome/peropsin, neuropsin and arthropsin. Our broader taxonomic sampling also allows us to infer three previously undescribed bilaterian paralogs, which we have named ‘xenopsins’, ‘bathyopsins’ and ‘chaopsins’. Because adding so many new bilaterian opsins changes the relationships between paralogs, we establish new, named hypotheses for these relationships, as often done in species-level phylogenetic analyses (see Table 1). In addition to new clade names, we also use Roman numerals for eumetazoan paralogs and Arabic numerals for bilaterian paralogs to help clarify which opsin clades are inferred as eumetazoan versus bilaterian paralogs at a glance in the text and figures.

**Figure 2.**
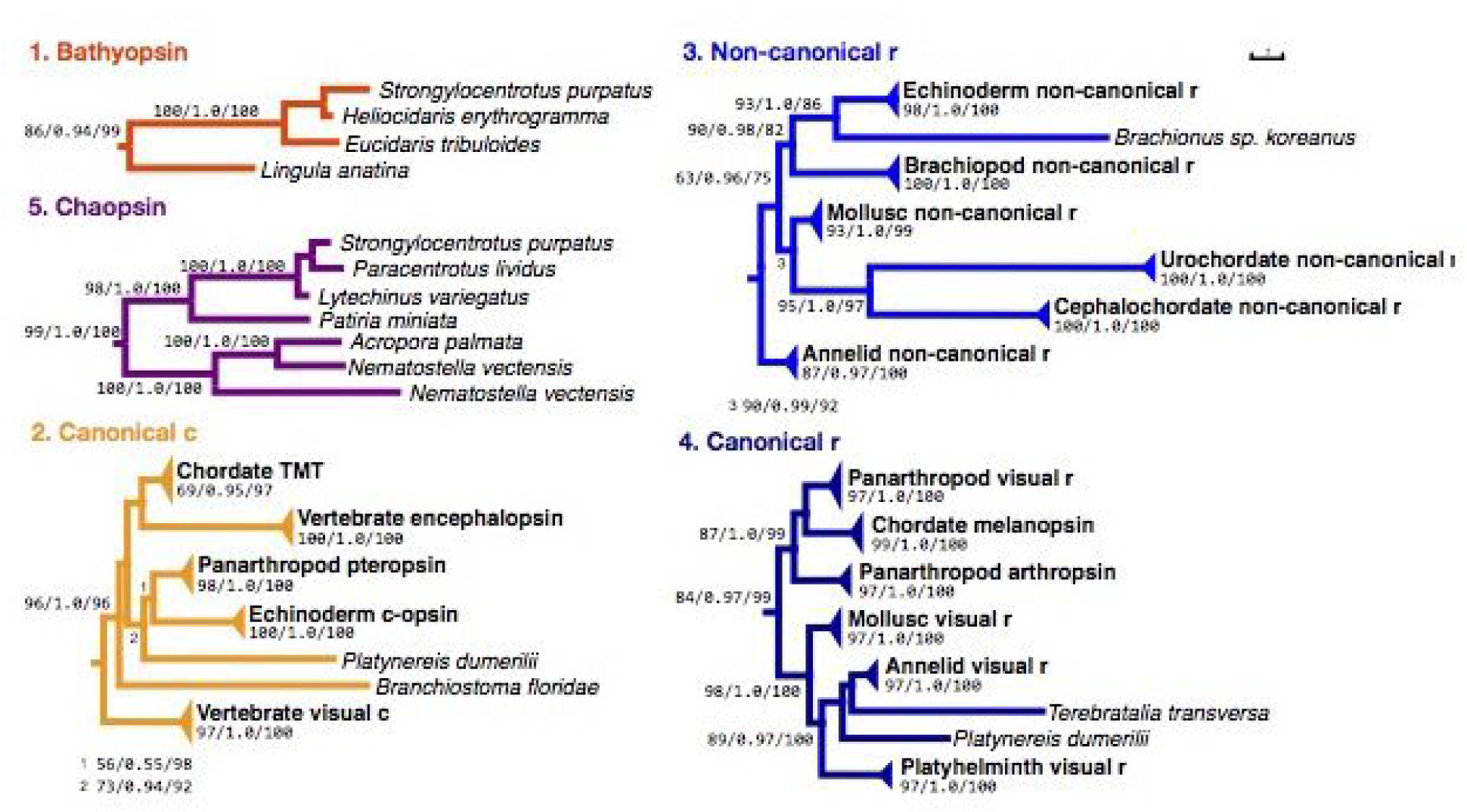
Opsin paralog trees for the tetraopsins and xenopsins, representing the relationships between opsins by phylum. Each tree shows opsin sequences collapsed by clade. Values below the clade name representSH-aLRT/aBayes/UFBoots. Only clades with bootstrap supports above 75% are shown. The full gene tree can be found in Supplemental Figure S1. Each asterisk ‘*’ on a branch represents a shortening by 5 branch length units.

**Table 1:**
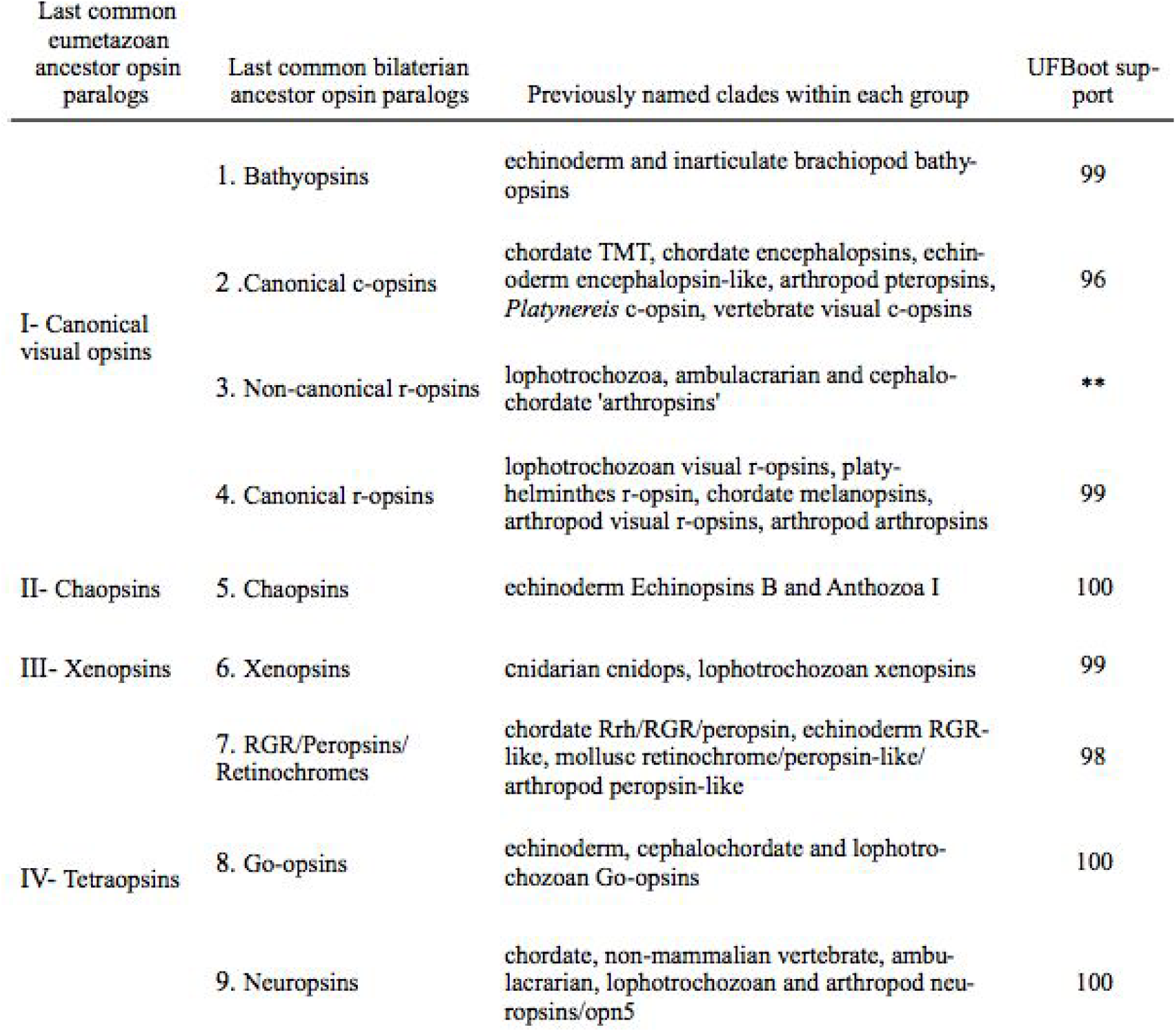
Summary of opsins present before bilaterians and present in the last common bilaterian ancestor. We considered UFBoot values above 95 as strong support for the monophyly of that group, but allowed branch rearrangements below that threshold. Asterisks (**) indicate support for the node based on reconciliation with the species tree.

### The last common eumetazoan ancestor had at least 4 opsin paralogs

All cnidarian opsins included in our analysis fell sister to three opsin paralogs also shared by bilaterians: the canonical visual opsins (I), chaopsins (II), and xenopsins (III) (see Figure 1 and Suppl. Figs S1 & S2). In our gene tree, Anthozoa II (Hering & Mayer 2014) is sister to the canonical c-opsins, but with mixed support from UFBoot (67) and single branch tests (aLRT = 91.2; aBayes = 0.998). Our reconciliation analysis minimizes duplications and losses, allowing rearrangement of nodes with low UFBoot values, and so places Anthozoa II sister to multiple bilaterian paralogs, including canonical c, non-canonical r, canonical r-opsins, and bathyopsins (Figure 1). However, given the difference in support for the placement of Anthozoa II based on bootstraps, single branch tests, and parsimony, we cannot confidently place these cnidarian opsins. On the other hand, the cnidarian Anthozoa I (Hering & Mayer 2014) are well supported as sister to echinoderm chaopsins (UFBoot = 100; aLRT = 98.6; aBayes = 1.0). Similarly, the group referred to as cnidops (Plachetzki et al. 2007) are also strongly supported as sister to lophotrochozoan sequences (UFBoot = 99; aLRT = 95.4; aBayes = 1.0), together comprising the xenopsins. Based on the positions of cnidarian opsins, we infer that these three opsin paralogs arose prior to the split of cnidarians + bilaterians, and were thus present in the last common ancestor of eumetazoans. Although we did not find extant cnidarian tetraopsins, we infer from our reconciled tree infers that the last common eumetazoan ancestor did have a tetraopsin, raising our estimate of eumetazoan opsin paralogs to at least four.

### Early bilaterian ancestors had at least 9 different opsin paralogs

#### I-Canonical visual opsins

This grouping consists of multiple clades of both previously and newly described opsins, encompassing the canonical visual opsins in both vertebrates and invertebrates. Because the relationships between these clades generally received low UFBoot support, their current placement together comes primarily by our reconciliation analysis to minimize duplication and loss by rearranging poorly supported nodes.

##### 1. New opsin group: Bathyopsins

The grou of opsin paralogs we have named bathyopsins is a small but well supported, monophyletic, bilaterian clade (see Figure 3, UFBoot=99). Sequences from the echinoderms, Echinopsins A (D’Aniello et al. 2015), represent deuterostomes, and sequences from the genome of the brachiopod *Lingula* represent prostostomes.

**Figure 3.**
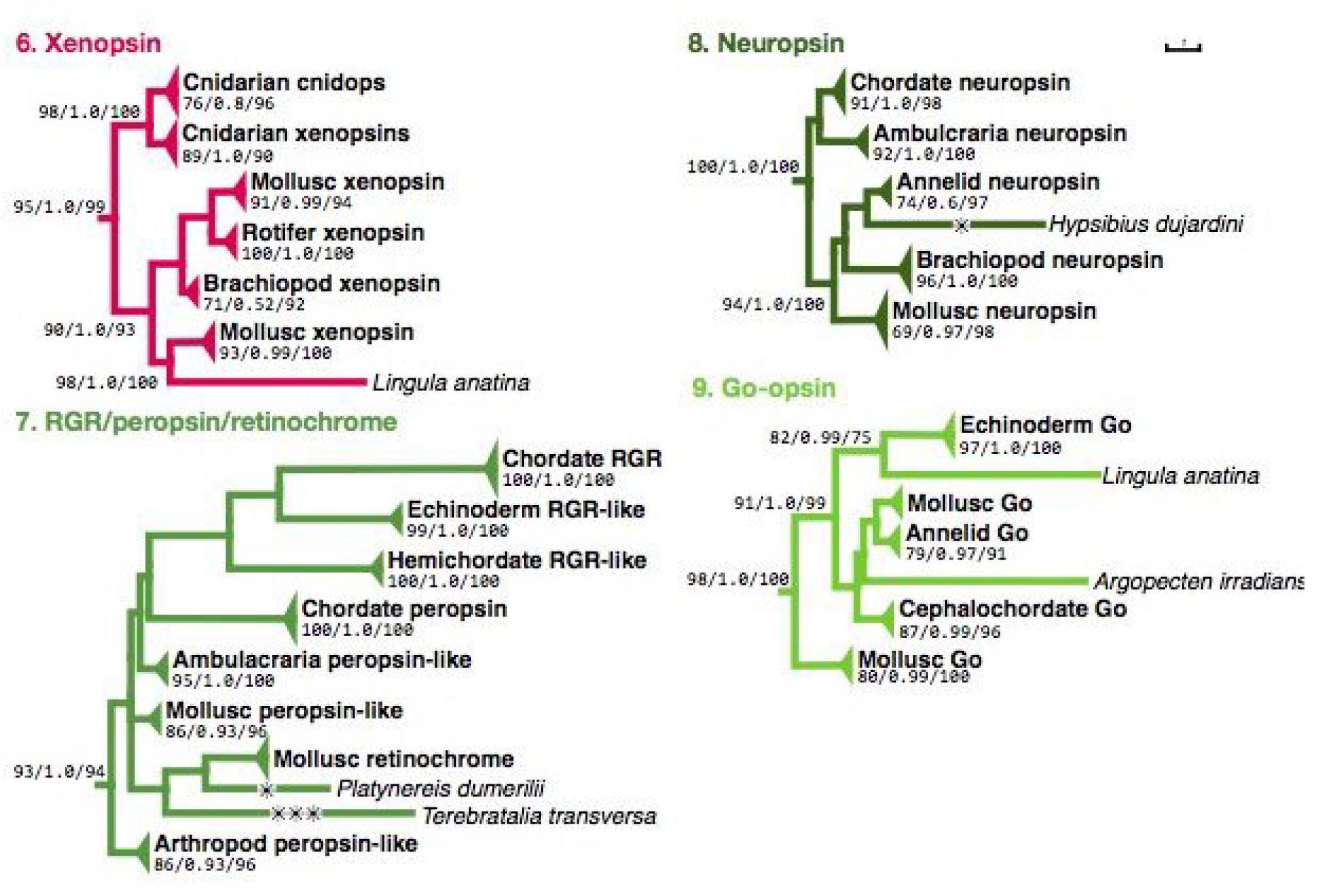
Opsin paralog trees for the Gqopsins and Copsins, representing the relationships between opsin orthologs by phylum. Each tree shows opsin sequences collapsed by clade. Values below the clade name represent SHaLRT/aBayes/UFBoots. Only clades with bootstrap supports above 75% are shown. The full gene tree can be found in Suppl. Figure S1.

**Figure 4.**
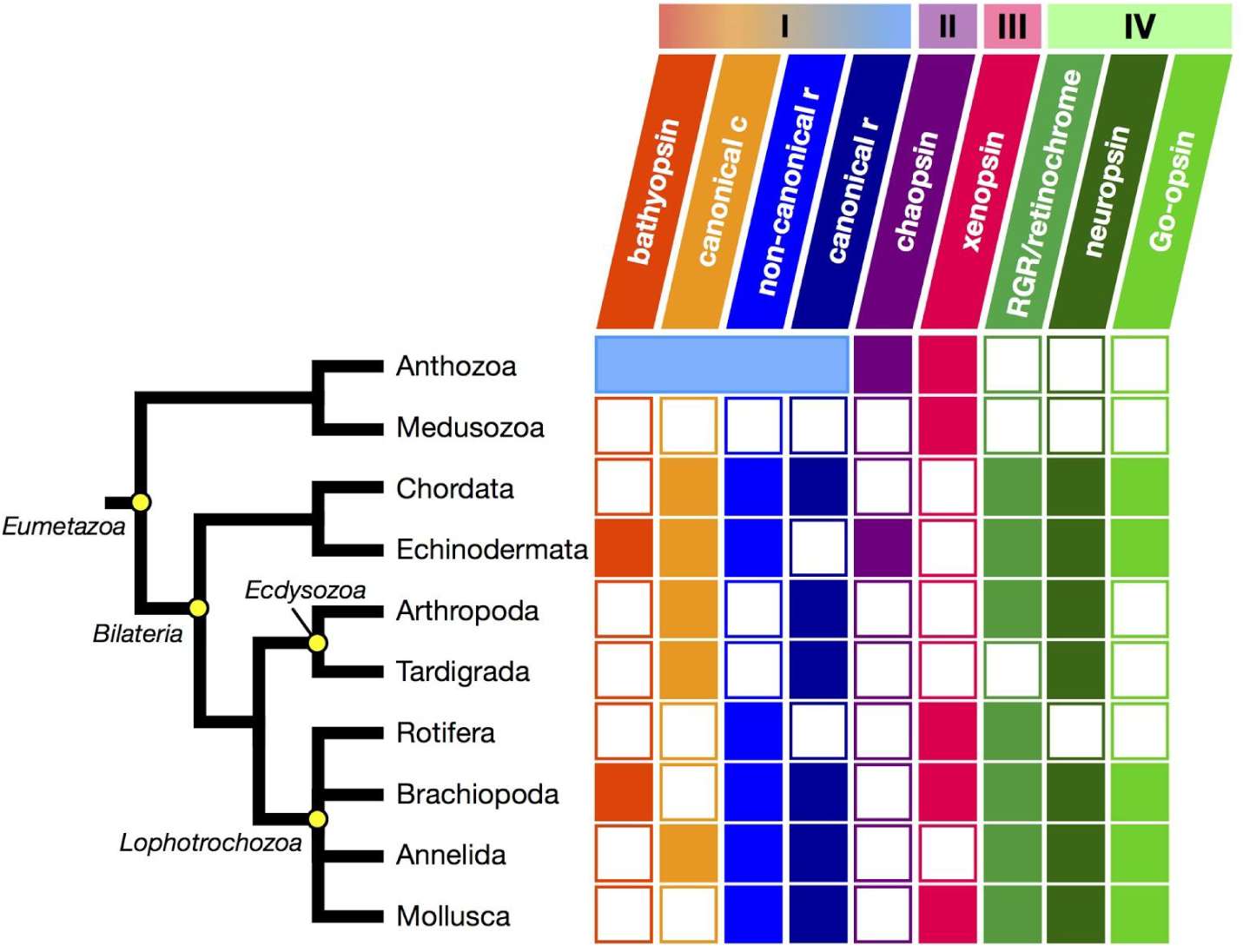
The history of opsins is marked by ancient diversity and subsequent losses of paralogs along different animal lineages. Summary of known opsin complements in major animal phyla. Major subdivisions of metazoans are indicated on the phylogeny as yellow dots with italic labels. Phyla are represented at the tips, except for cnidarians, which are broken down into the two major cnidarian splits. Colored bars with roman numerals indicate opsin paralogs present in the most recent ancestor of eumetazoans. The nine bilaterian opsin paralogs are indicated by slanted colored bars and full opsin names. Filled squares represent presence, empty squares absence of at least one sequence from the opsin paralog group for each phylum listed. Note that no extant phylum included in our analysis seems to have the full complement of bilaterian opsins. The maximum is 7 opsin paralogs in both echinoderms and brachiopods. The anthozoan Icanonical visual opsin paralog falls sister to bilaterian orthologs, and is indicated by the light blue bar.

**Figure 5.**
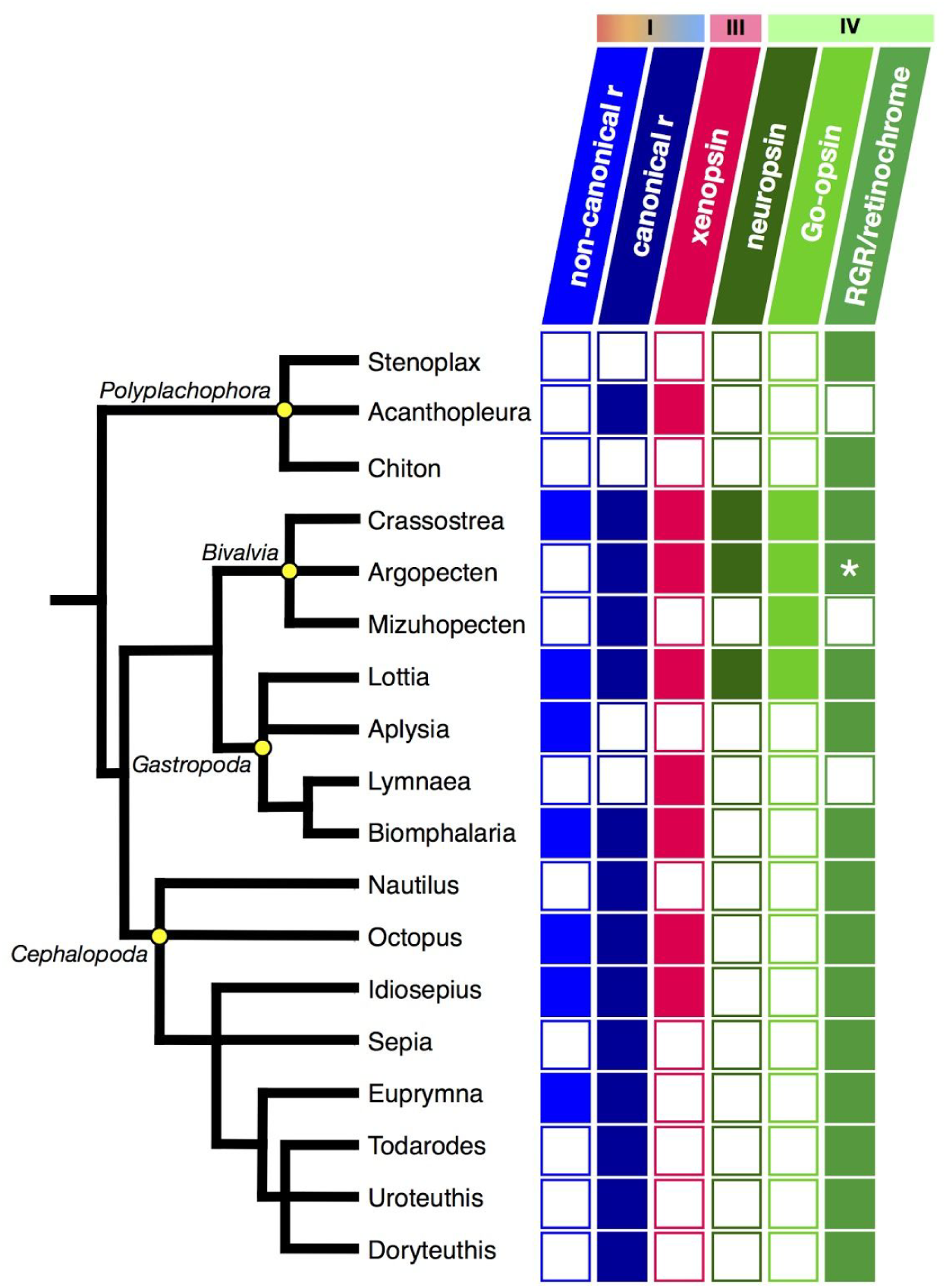
The ancestral mollusc likely had 7 opsins from six of the bilaterian paralog groups. Summary of known opsin complements within the molluscs. Colored bars with roman numerals indicate opsin paralogs present in the most recent ancestor of eumetazoans. The nine bilaterian opsin paralogs are indicated by slanted colored bars and full opsin names. Filled squares represent presence, empty squares absence of at least one sequence from the opsin paralog for each genus listed. The major classes of molluscs are noted with yellow dots and italic labels. *Argopecten irradians* retinochrome was not included our original analysis, but is present, noted by an asterisk (See Suppl. Figure S3 for RGR/retinochrome gene tree that includes this sequence).

##### 2. Canonical c-opsins

We have renamed as “canonical c-opsins” the monophyletic clade of bilaterian opsins such as vertebrate visual and brain c-opsins, arthropod pteropsins (Velarde et al. 2005), and *Platynereis* c-opsin (Arendt et al. 2004). We recovered the canonical c-opsins with high support (UFBoot=96, see Figure 3). Despite mining numerous mollusk transcriptomes for opsin sequences, we did not recover any additional lophotrochozoan or protostome c-opsins that clustered with the canonical c-opsins besides the single c-opsin reported from the annelid *Platynereis dumerilii.*

##### 3. Non-canonical r-opsins

The non-canonical r-opsins are sister to the canonical r-opsins with high support (UFBoot=100, see Figure 1), though we do not have strong support for the monophyly of the non-canonical r-opsins, even after reconciliation (see Figures 1 & 3; Suppl. Figure S1). The non-canonical r-opsins contain sequences from deuterostome lineages like echinoderms, hemichordates and cephalochordates, and previously unannotated sequences from protostomes groups that include annelids, brachiopods and molluscs.

##### 4. Canonical visual r-opsins

This opsin group is well supported (UFBoot=99, see Figure 3) and includes the following four clades: the canonical visual opsins of arthropods; the chordate melanopsins and arthropod arthroposins; the canonical visual opsins of mollusks; and the (presumably) visual r-opsins from annelids, brachiopods and platyhelminths.

#### II-5. New opsin group: chaopsins

The opsin group we have named chaopsins consists of two previously described clades of opsins, Anthozoa I (Hering & Mayer 2014) and the echinoderm Echinopsins B (D’Aniello et al. 2015). The grouping of these anthozoan and echinoderm sequences as monophyletic chaopsins is well supported (see Figure 3, UFBoot=100; aLRT=98.6; aBayes=1.0).

#### III-6. New opsin group: xenopsins

The opsin group we call xenopsins consists of sequences from a variety of lophotrochozoan protostomes (molluscs, rotifers and brachiopods) and cnidarian cnidops. This clade is well supported in our tree (see Figure 2, UFBoot=99; aLRT=95.4; aBayes=1.0). We did not find support for any other protostome (e.g. ecdysozoan) or deuterostome xenopsins. We recovered both the lophotrochozoan xenopsins and cnidarian cnidops with strong support (see Suppl Figure S1). The xenopsins are well supported as sister to the tetraopsins (UFBoot=98; aLRT=90.8; aBayes=1.0, see Figure 1 and Suppl. Figure 1)

Many xenopsins were initially described as c-opsin-like in previous analyses, including sequences from the genomes of the mollusks *Crassostrea gigas* and *Lottia gigantea* and the rotifer *Branchionus sp.*, and those from gene expression data generated from the larval eyes of the articulate brachiopod *Terebratalia transversa*, the optic lobes of *Octopus bimaculoides* and the adult eyes of *Idiosepius paradoxus* (Passamaneck et al. 2011; Albertin et al. 2015; Yoshida et al. 2015). However, we believe that the limited taxonomic scope of previous analyses lead to the incorrect classification of these sequences as c-opsin-like. Our tree is the first to include all of these sequences into a single analysis, and our results clearly support them as a monophyletic clade. Finally, in addition to xenopsins that were previously described, we found 7 new mollusc xenopsins from combing through transcriptomes (included in Suppl. Table S1).

#### IV-Tetraopsins

Similar to previous analyses (Porter et al. 2012; Hering & Mayer 2014; Feuda et al. 2014), we recover the tetraopsins (IV), formerly “RGR/Go” or “Group 4” opsins, as a monophyletic group with strong support (UFBoot = 100; aLRT = 98.9; aBayes = 1.0). They consist of RGR/retinochromes/peropsins, Go-opsins, and neuropsins. Because our tree shows strong support for these opsins as most closely related to each other, we have renamed this clade of opsins tetraopsins. Further, we find that each of the previously recognized major splits within tetraopsins has representatives from both protostomes and deuterostomes (see Figure 1).

##### 7. RGR/retinochromes/peropsins

The RGR/retinochrome/peropsin clade is well-supported by our tree (UFBoot=98, see Figure 2). Deuterostome RGRs include the original RGRs identified in vertebrates, as well as RGR-like sequences in cephalochordates, hemichordates, and echinoderms (Jiang et al. 1993; Holland et al. 2008; D’Aniello et al. 2015). Deuterostome peropsins include RRH from vertebrates as well as peropsin-like sequences from cephalochordates, hemichordates and echinoderms (Sun et al. 1997; Holland et al. 2008; D’Aniello et al. 2015). Protostome retinochromes include the originally described retinochromes from cephalopods, plus retinochrome-like sequences in bivalve and gastropod molluscs (Hara & Hara 1967; Katagiri et al. 2001). We recovered an additional 3 retinochrome-like sequences from mollusc transcriptomes, including 1 from the gastropod *Bithynia siamensis goniomphalos* and 2 from the chitons *Stenoplax conspicua* and *Chiton virgulatus.* In addition to the molluscs, we found retinochrome-like sequences in the brachiopod *Terebratalia transversa*, previously described as a Go-opsin (Passamaneck & Martindale 2013) and a sequence previously described as a peropsin in the annelid *Platynereis dumerilli* (Marlow et al. 2014). We also found a small clade of protostome sequences that fell outside of the protostome retinochromes, including 4 sequences from the genomes of the mollusks *Crassostrea gigas*, *Lottia gigantea* and *Octopus bimaculoides* (Albertin et al. 2015). Finally, non-insect arthropod peropsin-like sequences (Henze & Oakley 2015) also belonged in the clade of protostome retinochromes. It is unclear from our analysis whether RGR/retinochromes and peropsins are separate bilaterian paralogs. We did recover distinct groups, suggestive of of two bilaterian clades, but had low support values at these nodes, and so we collapsed the groups together (see Suppl. Figure S1).

##### 8. Neuropsins

The split between the protostome and deuterostome neuropsins is well supported (UFBoot=100, see Figure 2). Deuterostome neuropsins/opn5 sequences include a large clade of vertebrate and cephalochordate neuropsins, a large clade of non-mammalian vertebrate neuropsins, plus neuropsin-like sequences from the Ambulacraria (including those from both hemichordates and echinoderms). Neuropsins from protostomes include sequences from annelids, both *Platynereis dumerilli* (Gühmann et al. 2015) and *Capitella teleta* (Simakov et al. 2012), bivalve and gastropod molluscs, and from the brachiopod *Lingula anatina* (previously annotated as a peropsin). We recovered an additional bivalve neuropsin from the transcriptome of the scallop *Argopecten irradians.* We also found two pan-arthropod neuropsin-like sequences from water flea *Daphnia pulex* (Hering & Mayer 2014; Brandon 2015) and the tardigrade *Hypsibius dujardini* (Hering & Mayer 2014).

##### 9. Go-opsins

The deuterostome and protostome Go-opsins form a well supported clade of bilaterian opsins (UFBoot=100, see Figure 2). We recovered the same deuterostome Go-opsins from echinoderms and cephalochordates as identified from previous analyses (D’Aniello et al. 2015). From protostomes, we found previously described sequences of Go-opsins from both bivalve and gastropod mollusc, and also sequences from brachiopods and annelids. We also recovered a new Go-opsin from the transcriptome of the scallop *A. irradians.*

## Discussion

Reconstructing the evolutionary history of opsins is vital for understanding how light-detecting structures like eyes evolved. Unfortunately, the problem of how and when opsin diversity arose is made difficult by the large number of duplications and losses that have occurred within their evolutionary history. While most analyses of opsin diversity to date have focused on understanding the opsins present within a set of focal taxa (e.g. Plachetzki et al. 2007; Feuda et al. 2012; Hering & Mayer 2014; Feuda et al. 2014; D’Aniello et al. 2015), we included multiple poorly-sampled phyla to ensure the broadest phylogenetic scope to date, for a total of 248 species from 14 phyla. Our analysis reveals three previously unrecognized opsin paralogs in extant animals, and the surprising result that these three additional opsin paralogs likely arose early in the evolution of bilaterians, followed by losses and duplications within those opsins that remained.

Our first major finding is the presence of 9 bilaterian opsin paralogs in extant animals that we infer were also present in the last common ancestor of protostomes and deuterostomes. In addition to the 6 previously identified bilaterian opsins (c-opsin, r-opsin, melanopsin, Go-opsin, peropsin/RGR/retinochrome and neuropsin), we propose three additional bilaterian opsins— xenopsins, bathyopsins and chaopsins. While we acknowledge the need for additional sequence data to confirm the monophyly of these clades of opsin paralogs, our results are consistent with the hypothesis that these opsin paralogs were all present in the last common ancestor of Nephrozoa (protostomes + deuterostomes, excluding Xenacoelomorpha) and potentially bilaterians (depending on the opsins present in Xenacoelomorpha). Hints of the three new opsin groups we identified can be seen in previous opsin phylogenies (Hering & Mayer 2014; D’Aniello et al. 2015), but hypotheses for how these orphaned sequences relate to other better-studied opsins remained obscure with less broad taxonomic coverage.

For example, cnidarian cnidops have been difficult to place consistently within opsin phylogenies. We found that cnidops fall sister to the lophotrochozoan xenopsins with high bootstrap and branch support, suggesting they together form a monophyletic clade. Further, the hypothesis of xenopsins as the sister clade to the tetraopsins is also well supported both by UFBoot and single branch tests. If our reconciled gene tree is correct, the grouping of lophotrochozoan and cnidarian xenopsins suggests that xenopsins were present in both the bilaterian and eumetazoan ancestors. Our result differs from previous studies, (e.g. Plachetzki et al. 2007; Feuda et al. 2012; Bielecki et al. 2014), because these did not include bilaterian xenopsin-like sequences. Thus, Plachetzki et al. (2007) concluded that cnidops was its own eumetazoan opsin paralog, lost from bilaterians entirely. Both Feuda et al. (2012) and Bielecki et al. (2014) proposed cnidops as sister to the tetraopsins, and inferred that a tetraopsin was present in the eumetazoan ancestor. Our results are consistent with Feuda et al. (2012) and Bielecki et al. (2014), but our inferences differ— instead of the single eumetazoan tetraopsin, we infer two eumetazoan paralogs, xenopsins and tetraopsins.

Lophotrochozoan xenopsins are a well-supported monophyletic clade, suggesting that xenopsins were present in the lophotrochozoan ancestor. Interestingly, xenopsins are absent from publicly available *Platynereis* opsins and the *Capitella* and *Helobdella* genomes. However, because our sampling from annelids is so sparse given the large number of species in the phylum, it seems likely that annelid xenopsins could be uncovered after broader sampling. Xenopsins are also absent from both the ecdysozoan and deuterostome taxa included in our analysis. Given that arthropods, chordates, and echinoderms are now well-sampled for opsin diversity, it seems unlikely that xenopsins will ever be discovered in these phyla. Thus we hypothesize that the absence of xenopsins from these groups in our dataset reflects true losses of xenopsins from ecdysozoan and deuterostomes lineages. Given this hypothesis, we infer that xenopsins were lost at least three times in bilaterians: from ancestors of the annelids, Panarthropoda, and the deuterostomes.

Increased taxon sampling also allows us to hypothesize the bathyopsins and chaopsins as paralogs present in the last common ancestor of most bilaterians. These opsins are are unusual because of their extreme phylogenetic sparseness, suggesting that if our gene tree inference is correct, these opsin paralogs were lost in the majority of bilaterians. However, we interpret this sparseness as an indication that even our inclusive dataset may still be under-sampling true opsin diversity in animal phyla. Bathyopsins are found in only two phyla so far, Echinodermata and Brachiopoda, forming a well supported, monophyletic clade in our tree. Given that bathyopsins are represented by one deuterostome and one protostome, we infer that bathyopsins were present in the last common bilaterian ancestor. We have not found chordate or hemichordate representatives. In protostomes, we infer that the lophotrochozoan ancestor had bathyopsins, but since bathyopsins are unknown in ecdysozoa entirely, it is possible that they were lost in ecdysozoa after the lophotrochozoan/ecdysozoan split. Because opsins from chordates and arthropods are well sampled, it is unlikely that these phyla possess bathyopsins. We have not uncovered annelid, mollusc or rotifer bathyopsins, yet sparse sampling in these taxa makes it possible that opsin surveys from lophotrochozoans will reveal additional members of the bathyopsins in these phyla.

We found chaopsins in only two phyla so far, echinoderms and cnidarians, and their monophyly is supported by both high UFboots and single branch tests. Given our data set and analysis, we hypothesize that chaopsins were lost up to three times in bilaterians: twice from deuterostomes (chordates and hemichordates) and once in the ancestor of all protostomes (including both ecdysozoans and lophotrochozoans). We also find that anthozoans are the only cnidarians that have chaopsins, which suggests another potential loss of chaopsins from the ancestor of hydrozoans and cubozoans. As with the other new opsin types we have described, we are more confident that chaopsins are truly lost from chordates and arthropods compared to the undersampled lophotrochozoans.

Our second major finding is the eumetazoan ancestor likely had at least four opsin paralogs, based on the distribution of cnidarian opsins in our analyses. Instead of inferring eumetazoan c-, r-, and tetraopsins as previously reported (Feuda et al. 2012; Bielecki et al. 2014; Hering & Mayer 2014), we found cnidarian orthologs of the canonical visual opsins, xenopsins and chaopsins. Along with these three eumetazoan opsins, we infer that the last common eumetazoan ancestor also had a tetraopsin, but this paralog is not present in the cnidarians we surveyed. However, because cnidarians are not well sampled, it is possible that a cnidarian ortholog of the bilaterian tetraopsins may be uncovered. Overall, we successfully identified well-supported bilaterian orthologs of at least two cnidarian opsins— cnidops as xenopsins, and Anthozoa II opsins as chaopsins, and infer that the last common ancestor of eumetazoans must have had at least 4 different opsins. Adding more opsins from Cnidaria, Ctenophora, and Xenacoelomorpha may help solidify deeper relationships between well documented opsin paralogs like the canonical c-and r-opsins and the opsin paralogs we have identified in this analysis.

We used a traditional animal phylogeny to reconcile our gene tree, with ctenophores placed sister to cnidarians, and these two phyla together as sister to bilaterians (“ctenophore-in” hypothesis). This traditional view is recently challenged by multiple studies that instead place ctenophores sister to all other animals (“ctenophore-out” hypothesis) (Dunn et al. 2008; Ryan et al. 2013; Moroz et al. 2014; Borowiec et al. 2015; Pisani et al. 2015; Halanych et al. 2016). There seem to be two opsin paralogs in ctenophores, but the relationship between those opsin paralogs and opsins from other animals is contentious, particularly the placement of Mnemiopsis 3 (Feuda et al. 2014; Schnitzler et al. 2012). Although Mnemiopsis 3 does have the conserved lysine that aligns at bovine rhodopsin position 296, it was excluded from (Hering & Mayer 2014) because there is an insert that is absent from the other *Mnemiopsis* opsins. Its placement in the metazoan opsin phylogeny is also highly sensitive to outgroups as seen in (Schnitzler et al. 2012; Feuda et al. 2014). For these reasons, we did not include Mnemiopsis 3 in our analysis. Overall, our estimates of opsin repertoires in the last common eumetazoan ancestor and early bilaterians are likely not greatly impacted by the current controversy about the relationship between ctenophores and other animals (Borowiec et al. 2015; Pisani et al. 2015; Halanych et al. 2016; Pisani et al. 2016). Ctenophores-in might suggest that they have lost some of the eumetazoan opsin paralogs, while ctenophores-out suggests a very early origin of the first opsin, and a likely loss of opsins in sponges. At present our results do not distinguish between ctenophore-in and ctenophore-out, as the ctenophore opsins we included were not placed in the animal opsin phylogeny with high support.

Opsin evolution is surprisingly complex, hinting at just how much we have yet to learn about how animals use opsins, how these functions shaped the evolution of the gene family, and the physiology and behaviors that require opsins. Besides spatial vision, opsins are used for myriad purposes, e.g. as depth-gauges or for circadian rhythms (Bennett 1979; Lythgoe 1979; Bybee et al. 2012). Further, opsins are not only expressed in eyes, but also across the bodies of animals (reviewed in Ramirez et al. 2011). It is not yet clear to what extent the loss of an opsin paralog within an animal lineage suggests the concomitant loss of the organismal function, or whether other opsin paralogs can take over that function. At present, we have no functional data for the majority of the 700+ opsins included in this analysis, but current data suggest different animal phyla use related opsins for different purposes. For example, r-opsins likely mediate vision in many protostome eyes, but the related orthologous melanopsins in vertebrate retinal ganglion cells only have roles in non-visual tasks. While opsins are canonical light detectors, two recent studies have shown roles for opsins in both heat sensing and detecting mechanical stimuli in *Drosophila* (Shen et al. 2011; Senthilan et al. 2012). These studies provide a tantalizing glimpse into opsin functions in sensory modalities besides light detection. Without understanding the true extent of opsin diversity, we cannot understand opsin evolution, the evolution of eyes and other light sensors, or even how a complex trait like eyes can evolve.

## Acknowledgments

This work was supported by the National Science Foundation [grant numbers IOS-1457148 to D.I.S & T.H.O.; DMR-1121053, CNS-0960316 to the California NanoSystems Institute and Materials Research Laboratory through the Center for Scientific Computing]. We would like to thank Davide Pisani and Roberto Feuda for comments on an early version of this manuscript, and the anonymous reviewers on our first submission.

## Data deposition in NCBI

KX550901

KX550902

KX550903

KX550904

KX550905

KX550906

KX550907

KX714605

KX714606

KX714607

KX714608

